# The lung extracellular matrix protein landscape in severe early-onset and moderate chronic obstructive pulmonary disease

**DOI:** 10.1101/2023.10.20.562391

**Authors:** M.M. Joglekar, N.J. Bekker, M.L. Koloko Ngassie, J.M. Vonk, T. Borghuis, M.A. Reinders-Luinge, J. Bakker, R.R. Woldhuis, S.D. Pouwels, B.N. Melgert, W. Timens, C.A. Brandsma, J.K. Burgess

## Abstract

Extracellular matrix (ECM) remodeling has been implicated in the irreversible obstruction of airways and destruction of alveolar tissue in chronic obstructive pulmonary disease (COPD). Studies investigating differences in the lung ECM in COPD have mainly focused on some collagens and elastin, leaving an array of ECM components unexplored. We investigated the differences in the ECM landscape comparing severe-early onset (SEO-) COPD and moderate COPD to control lung tissue for collagen type I α chain 1 (COL1A1), COL6A1, COL6A2, COL14A1, fibulin 2 and 5 (FBLN2, FBLN5), latent transforming growth factor-beta binding protein 4 (LTBP4), lumican (LUM), versican (VCAN), decorin (DCN), and elastin (ELN) using image analysis and statistical modelling. Percentage area and/or mean intensity of expression of LUM in the parenchyma, and COL1A1, FBLN2, LTBP4, DCN, and VCAN in the airway walls, was proportionally lower in COPD compared to controls. Lowered levels of most ECM proteins were associated with decreasing FEV_1_ measurements, indicating a relationship with disease severity. Furthermore, we identified six unique ECM signatures where LUM and COL6A1 in parenchyma and COL1A1, FBLN5, DCN, and VCAN in airway walls appear essential in reflecting the presence and severity of COPD. These signatures emphasize the need to examine groups of proteins to represent an overall difference in the ECM landscape in COPD, that are more likely to be related to functional effects, than individual proteins. Our study revealed differences in the lung ECM landscape between control and COPD and between SEO and moderate COPD signifying distinct pathological processes in the different subgroups.

**NEW & NOTEWORTHY:** Our study identified COPD-associated differences in the lung ECM composition. We highlight the compartmental differences in the ECM landscape in different subtypes of COPD. The most prominent differences were observed for severe-early onset COPD. Moreover, we identified unique ECM signatures that describe airway walls and parenchyma providing insight into the intertwined nature and complexity of ECM changes in COPD that together drive ECM remodeling and may contribute to disease pathogenesis.

## 1. INTRODUCTION

Chronic obstructive pulmonary disease (COPD) is a lung disease with increasing prevalence globally (1). The pathogenesis of COPD has been largely attributed to prolonged exposure to cigarette smoke, air pollution, or occupational pollutants in combination with genetic predisposition. These factors initiate a process that causes a decline in lung function which is evaluated clinically by measuring the forced expiratory volume in 1 second (FEV_1_) (2). Based on FEV_1_ measurements, the Global Initiative for Chronic Obstructive Lung Diseases (GOLD) has classified COPD into four stages-COPD stage I, II, III and IV (3).

COPD patients present with several phenotypes, reflecting the heterogeneity of the disease, including chronic bronchitis in the airways and emphysema in the alveoli distal to terminal bronchioles. Deposition and remodeling of the extracellular matrix (ECM) in the airway wall contributes to irreversible airway wall thickening. Conversely, emphysema is characterized by the destruction of alveolar tissue causing loosening and ultimately loss of alveolar attachments and elastic recoil. The ECM is a three-dimensional network of proteins, proteoglycans, and glycosaminoglycans that provides structural support, including tensile strength and elasticity to the lung and essential biochemical and biophysical cues to cells (4). Several studies, often with conflicting results, have highlighted differences in ECM content in COPD as previously reviewed (5, 6).

Several studies have drawn parallels between physiological lung aging and COPD, as these processes share multiple hallmarks including dysregulated ECM remodeling. Thus, “accelerated aging” is often considered one of the main features of COPD (7–9). Apart from the most common phenotypes of COPD (bronchitis and emphysema), the severity and age of onset of the disease can define certain subgroups of patients. Historically, COPD is considered a disease of the elderly, predominantly male smokers. However, it is now clear that a subgroup of patients, with a high prevalence among women, develop very severe COPD at a much earlier age (often younger than 55 years) and this group is referred to as severe early onset (SEO)-COPD patients (8, 10, 11).

Our group recently reported age-associated ECM differences in human lung tissue using a combination of transcriptomic and proteomic analyzes (12). Seven ECM and ECM-associated proteins including collagen type I α chain 1 (COL1A1), COL6A1, COL6A2, COL14A1, fibulin 2 (FBLN2), latent transforming growth factor β binding protein 4 (LTBP4), and lumican (LUM) were found to have higher expression with increasing age at gene and protein levels in healthy subjects. Immunohistochemical studies further illustrated higher levels of COL1A1, COL6A2, COL14A1, and LUM in different lung compartments with age. We hypothesized that the proteins involved in the aging processes of the lung also play a key role in the pathology of COPD. In the present study, we aimed to investigate whether these seven age-related proteins showed similar differences in COPD lung tissue and whether the degree of these differences was related to disease severity (i.e. FEV1) and COPD subgroups, i.e. SEO-COPD and moderate COPD. Additionally, other important matrix proteins including fibulin 5 (FBLN5), decorin (DCN), versican (VCAN), and elastin (ELN) that are of known biological relevance in the pathology of COPD (13–15) were also investigated to identify differences in the ECM landscape between control, SEO-COPD, and moderate COPD lungs. Finally, statistical modeling was performed on the combined set of ECM and ECM-associated proteins to define COPD ECM signatures in the different compartments of the lung, specific for COPD, SEO-COPD and moderate COPD.

## 2. MATERIALS AND METHODS

### 2.1 Ethics statements

The study was conducted in accordance to the Research Code of the University Medical Center Groningen (UMCG), as stated on https://umcgresearch.org/w/research-code-umcg as well as national ethical and professional guidelines Code of Conduct for Health Research (https://www.coreon.org/wp-content/uploads/2023/06/Code-of-Conduct-for-Health-Research-2022.pdf). The use of left-over lung tissue in this study was not subject to Medical Research Human Subjects Act in the Netherlands, as confirmed by a statement of the Medical Ethical Committee of the University Medical Center Groningen and therefore exempt from consent according to national laws (Dutch laws: Medical Treatment Agreement Act (WGBO) art 458 / GDPR art 9/ UAVG art 24). All donor material and clinical information were deidentified prior to experimental procedures, blinding any identifiable information to the investigators.

### 2.2 Subjects

COPD and control human lung tissues were obtained from material leftover following lung transplants and tumor resection surgeries at the University Medical Center Groningen (Groningen, The Netherlands). In the latter, only lung tissue sections distant from the resected tumor, that appeared normal upon macroscopic and histological evaluation were accepted for use. This study was part of the HOLLAND (HistopathOlogy of Lung Aging aNd COPD) cohort (12). The donors were selected based on the following inclusion criteria:

a. SEO-COPD patients: FEV_1_%pred <40%, FEV_1_/FVC <70%, and age ≤55years at the time of lung transplant surgery, ex-smokers (10, 11, 16, 17).
b. Non-COPD control subjects (matched with SEO-COPD): FEV_1_/FVC >70%, age <65 years at the time of surgery, ex-smokers.
c. Moderate COPD patients: FEV_1_%pred 40-80%, FEV_1_/FVC <70%, age >65 years at the time of surgery, ex-smokers.
d. Non-COPD control subjects (matched with moderate COPD): FEV_1_/FVC >70%, age >65 years at the time of surgery, ex-smokers.

For patients where both pre- and post-bronchodilator FEV_1_, FVC and FEV_1_/FVC measurements were available, the best measurement was chosen for inclusion.

### 2.3 Immunohistochemistry

Briefly, lung tissue obtained from COPD (n= 26) and control (n=18) donors was formalin-fixed paraffin-embedded (FFPE) and cut into 6µM sections. Prior to staining, the sections were deparaffinized and rehydrated. The full methodology for staining for COL1A1, COL6A1, COL6A2, COL14A1, FBLN2, LTBP4, and LUM has been previously described (12). For FBLN5, VCAN, ELN, and DCN sections were treated with Tris/EDTA buffer (10mM, pH 9) for antigen retrieval. The sections were washed with PBS and endogenous peroxidase activity was blocked using hydrogen peroxidase (0.3%) for 30min at room temperature. Following PBS washes, primary antibodies diluted in 1% BSA/PBS for FBLN5 (1:8000, Mouse Anti-Fibulin 5/DANCE Antibody 1G6A4, Novus Biologicals), DCN (1:1500, Mouse Anti-Dermatan Sulfate Proteoglycan Antibody 6B6, Seikagaku), VCAN (1:200, Mouse Anti-Versican Antibody 2B1, Seikagaku), and ELN (1:400, Rabbit Anti-Elastin Antibody CL55011AP, Cedarlane Labs) were added to the respective sections for 1 hour at room temperature. After the incubation period, the sections were washed and horseradish peroxidase conjugated secondary antibody Rabbit Anti-Mouse (1:100, P0260, Dako, Denmark) was added to FBLN2, DCN, and VCAN staining and Goat Anti-Rabbit (1:100, P0448, Dako, Denmark) was added to ELN sections in 1% BSA-PBS containing 1% human serum. Similarly, tertiary antibody Goat Anti-Rabbit (1:100, P0448, Dako, Denmark) was used to stain FBLN5, DCN, and VCAN sections while Rabbit Anti-Goat (1:100, P0449, Dako, Denmark) was used for ELN in 1% BSA-PBS containing 1% human serum for after washing away the secondary antibodies. Negative controls (no primary antibody) were also included. Finally, positive staining in the sections was visualized with Vector® NovaRED® substrate (SK-4800, Vector Laboratories, Canada). Haematoxylin was used to counterstain these sections. All sections used for examination of a given protein were stained at the same time. The sections were dehydrated and mounted and scanned at 40x using a digital slide scanner Hamamatsu Nanozoomer 2.0HT (Hamamatsu Photonic K.K., Japan). Aperio ImageScope (v12.4.6.5003, Leica Biosystems, Germany) was used to view these digital images.

### 2.4 Image Analysis

The expression and localization of proteins in COPD compared to control donors was investigated in the airways and parenchymal regions of the lung tissue, as described previously (12). Individual compartments were extracted from tissue scans using Aperio ImageScope. Depending on the donor, up to a maximum of 10 airways (<2mm in diameter) were extracted from each section. After extraction, specific areas of interest including parenchyma and airway walls were further isolated using Adobe Photoshop 2023 (Adobe Inc. California, USA). Fiji (ImageJ) (18) was used to quantify percentage area stained (relative to the amount of tissue present on the slide) and mean intensity of pixels reaching the threshold for positive staining in the parenchymal regions and airway walls separately using color deconvolution plugin developed by Landini *et al.* (19) and as described in detailed by Koloko Ngassie *et al.* (12). Data sorting and calculations were performed using R software (version 4.2.3, USA).

### 2.5 Statistical Analysis

Donor characteristics including age, sex, pack-years, FEV_1_%pred, and FEV_1_/FVC were compared between subgroups of COPD and matched controls group using Mann-Whitney U tests. Percentage area and mean intensity of proteins that were not normally distributed were log (natural) transformed, and these transformed values were used for further analyzes. The differences in percentage area and mean intensity of protein levels in COPD and control lung tissue were examined using linear regression analysis for parenchyma and linear mixed effects regression analysis with a random effect on intercept per subject for airway walls. These models were also used to compare sub-groups of COPD (moderate COPD and SEO-COPD) to their respective matched controls. Regression coefficients with the 95% confidence intervals were plotted. Furthermore, linear and linear mixed models were also used to investigate associations between FEV_1_%pred measurements and percentage area or mean intensity of each protein in parenchyma and airway walls respectively.

Principal component analysis (PCA) with varimax rotation was used to identify unique ECM and ECM-associated protein signatures in COPD. Z scores of raw protein values (non-transformed or log transformed) were used as input and ‘n’ components, were extracted that cumulatively explained at least 80% of the variance. The component scores were saved for further analyzes. Component scores of COPD and control donors were compared per component using linear and linear mixed modelling for parenchyma and airway wall respectively. The comparisons of component scores between subgroups of COPD (SEO-COPD or moderate COPD) to all controls were performed using linear and linear mixed modelling, corrected for age. A p-value of <0.05 was considered significant. Data were analyzed using IBM SPSS version 28.0.1.0(142). Scatter and forest plots were created in GraphPad Prism version 8.0.0.

## 3. RESULTS

### 3.1 Patient characteristics

The clinical parameters of donors included in this study are summarized in **Table 1**. All subjects included in this study were ex-smokers. The COPD group (n=26) was comprised of moderate COPD (n=14) and SEO-COPD (n=12). For analyzes that investigated the effect of COPD subgroups, the control group (n=18) was divided into older control (n=9) subjects matched in age and sex to moderate COPD and younger control subjects (n=9) matched in age and sex to SEO-COPD (n=9).

**Table 1:** Characteristics of donors included in this study. SEO-COPD (n=12) and moderate COPD (n=14) were matched in terms of age, sex, and smoking status to respective younger and older control groups. A p value of <0.05 was considered significant.

|  | Younger controls | SEO-COPD | Older controls | Moderate COPD |
| --- | --- | --- | --- | --- |
| **Age in years**<br>**(Median, range)** | 55.0 (43.0- 62.0) | 51.5 (47.0- 55.0) | 74.0 (67.0-81.0) | 72.0 (67.0- 81.0) |
| **Sex**<br>**(Female/Male)** | 6/3 | 8/4 | 3/6 | 2/12 |
| **Pack Years**<br>**(Median, range)** | 15.0 (1.5-40.0) | 25.0 (6.0-54.0) | 35.0 (5.0-52.5) | 41.0 (10.0-65.0) |
| **FEV<sub>1</sub>%pred**<br>**(Median %**<br>**predicted, range)** | 103.6 (85.0-127.0) | 19.3 (14.9- 23.6)<sup>§</sup> | 91.5 (75.8-133.0) | 62.0 (49.7-75.4)<sup>&</sup> |
| **FEV<sub>1</sub>/FVC**<br>**(Median, range)** | 76.0 (69.0<sup>^</sup>-86.6) | 27.3 (20.5- 68.0)<sup>§</sup> | 72.0 (69.0- 86.1) | 60.8 (43.6- 67.9)<sup>&</sup> |

FEV<sub>1</sub>%pred: forced expiratory volume in 1 second % predicted; FVC: forced vital capacity; SEO-COPD: Severe early-onset COPD patients. <sup>^</sup>Two controls had an FEV<sub>1</sub>/FVC of 69% due to a relatively high FVC in combination with normal FEV<sub>1</sub>. <sup>§</sup>p<0.0005 and <sup>&</sup>p<0.0005 indicates the comparison between SEO-COPD and younger controls and moderate COPD and older controls, respectively.

### 3.2 Localization of ECM and ECM-associated proteins in lung tissue

The immunohistochemically stained lung tissues were examined for the localization of proteins in COPD and control lung tissue. Some examples of staining of COL1A1, COL6A1, COL6A2, COL14A1, FBLN2, FBLN5, LTBP4, LUM, DCN, VCAN, and ELN for each group are depicted in **Figure 1**. The localization of each staining has been summarized in **Table 2**. Briefly, all eleven proteins were detected in the parenchyma and COL14A1, LTBP4, and LUM staining was also detected in the epithelial layer. Within the airway wall, all proteins were detected in the airway adventitia and submucosa apart from DCN which was donor dependent. In the vicinity of the airway smooth muscle, COL6A1, COL6A2, COL14A1, LTBP4, and LUM were detected along with COL1A1 in some donors.

**Table 2:** Localization of ECM and ECM-associated proteins in lung tissue. An overview of positively stained areas in lung tissue.

| ECM protein | Parenchyma | Epithelial layer | Airway wall |  |  |
| --- | --- | --- | --- | --- | --- |
|  |  |  | SM | AA | ASM |
| COL1A1 | + | - | + | + | -/+ |
| COL6A1 | + | - | + | + | + |
| COL6A2 | + | - | + | + | + |
| COL14A1 | + | + | + | + | + |
| FBLN2 | + | - | + | + | - |
| FBLN5 | + | - | + | + | - |
| LTBP4 | + | + | + | + | + |
| LUM | + | + | + | + | + |
| DCN | + | - | -/+ | + | - |
| VCAN | + | - | + | + | - |
| ELN | + | - | + | + | - |

ECM: extracellular matrix; COL1A1: collagen type I $\alpha$ chain 1; COL6A1: collagen type VI $\alpha$ chain 1; COL6A2: collagen type VI $\alpha$ chain 2; COL14A1: collagen type XIV $\alpha$ chain 1; FBLN2: fibulin 2; FBLN5: fibulin 5; LTBP4: latent transforming growth factor binding protein 4; LUM: lumican; DCN: decorin; VCAN: versican; ELN: elastin; SM: submucosa; AA: airway adventitia; ASM: airway smooth muscle, (-) no staining, (+): positive staining.

**Figure 1:**
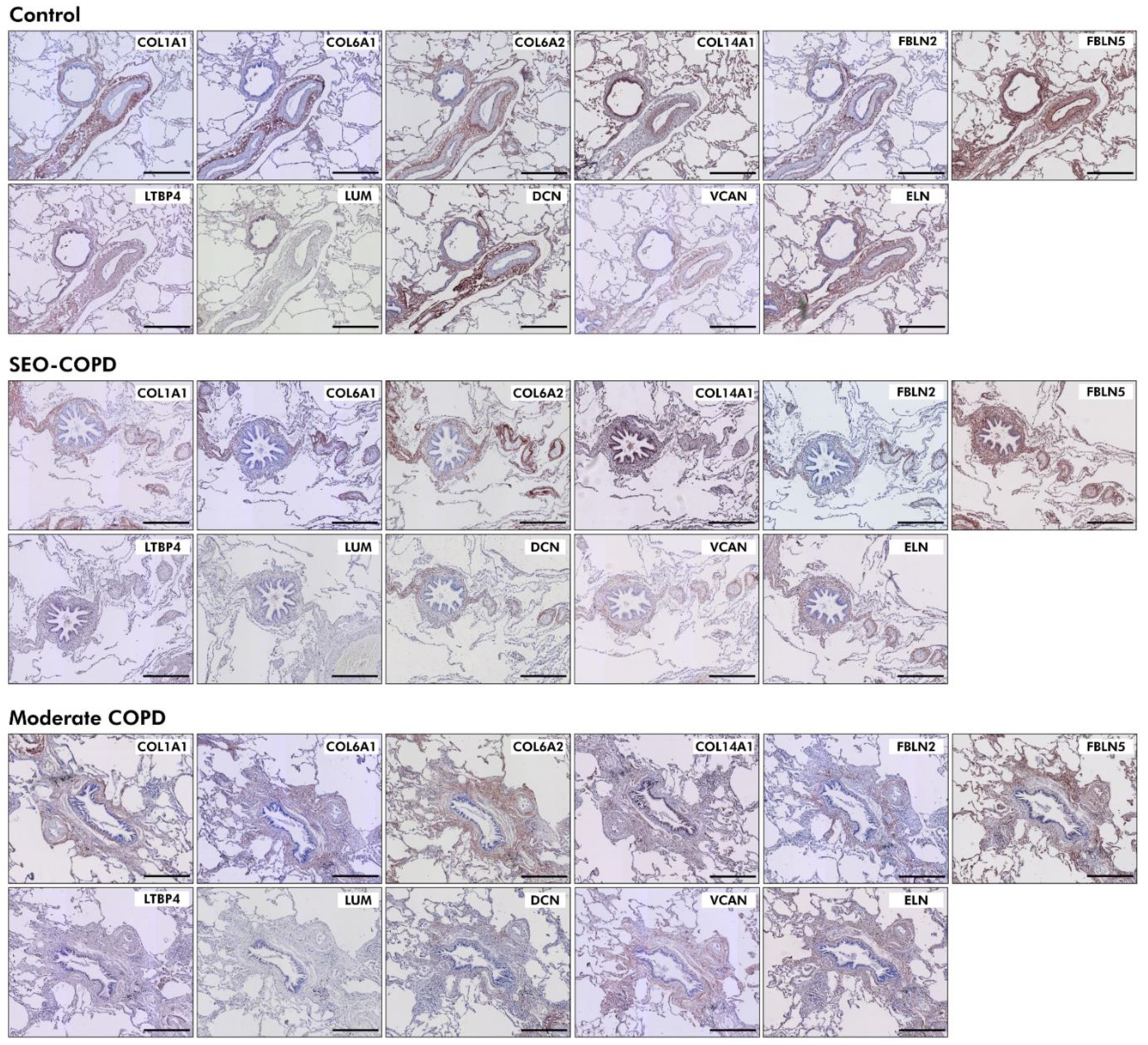
Localization of ECM and ECM-associated proteins in control, SEO and moderate COPD parenchyma and airway walls. FFPE lung tissue sections from control and COPD donors were stained using immunohistochemistry for ECM and ECM-associated proteins with specific signals being detected with Nova red (red) and counterstained with hematoxylin (blue). Sections were scanned at 40x magnification using a digital slide scanner. Scale bar = 400um. Images are representative of protein detection patterns seen in control donors (n=18), SEO-COPD (n=12), and moderate COPD (n=14). ECM: extracellular matrix; SEO-COPD: Severe early-onset COPD patients; FFPE: formalin fixed paraffin embedded; COL1A1: collagen type I α chain 1; COL6A1: collagen type VI α chain 1; COL6A2: collagen type VI α chain 2; COL14A1: collagen type XIV α chain 1; FBLN2: fibulin 2; FBLN5: fibulin 5; LTBP4: latent transforming growth factor binding protein 4; LUM: lumican; DCN: decorin; VCAN: versican; ELN: elastin.

### 3.3 Distribution of ECM and ECM-associated proteins in COPD lung tissue compared to matched controls

The distribution of ECM and ECM-associated proteins that have been previously associated with age related changes, or with COPD pathology, were investigated using image analysis and statistical modeling. Percentage area and mean intensity of positive pixels for each ECM protein was compared between COPD and matched controls, followed by subgroup analysis for SEO and moderate COPD compared to their matched controls.

#### 3.3.1 Differences in lung parenchyma; Lower proportional levels of LUM in COPD and SEO-COPD

When comparing staining of ECM and ECM-associated proteins in lung parenchyma between all COPD patients and all controls, we observed a lower proportional percentage area of the tissue that was positive for LUM (p=0.010) and a lower mean intensity of the positive LUM pixels (p=0.022) in COPD compared to control tissue (**Figure 2A, D**). Subgroup analysis demonstrated that this lower LUM percentage area was most apparent in SEO-COPD (p=0.012) (**Figure 2B**). There were no significant differences observed for the other ECM proteins between COPD and control, nor in the SEO or moderate COPD subgroup analysis. (**Figure 2C, E, F**). The percentage area or mean intensity of expression of some ECM proteins was significantly associated with age as also demonstrated in our previous publication (12). However, this did not affect the differences between COPD and control because these cohorts were age-matched (data not shown).

**Figure 2:**
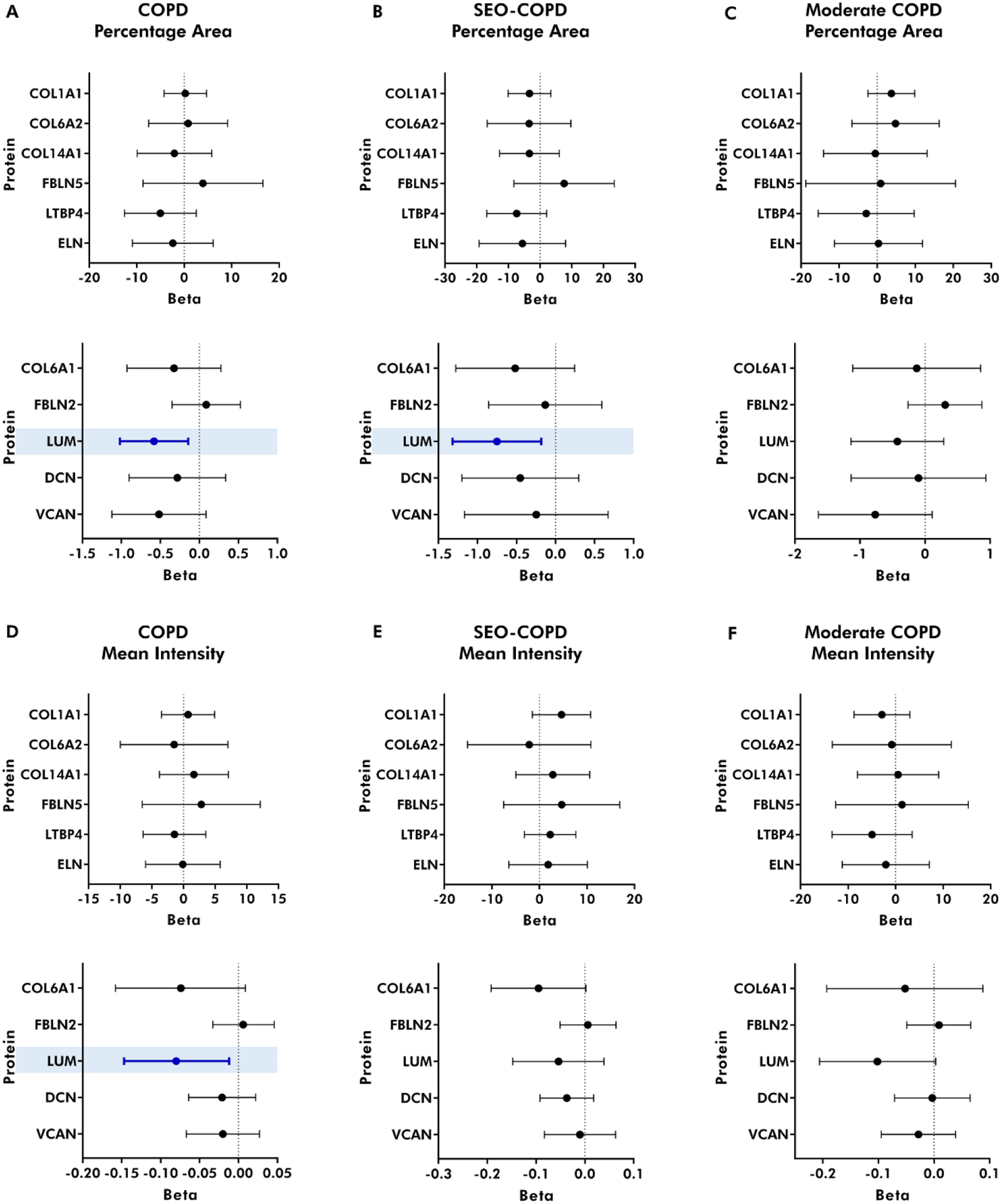
Forest plots of regression coefficients for percentage area and mean intensity of ECM and ECM-associated proteins in the parenchyma of COPD, SEO-COPD, and moderate COPD compared to their respective control groups. ECM proteins in FFPE tissue sections from controls (n=18), SEO-COPD (n=12), and moderate COPD (n=14) were detected using immunohistochemistry. Parenchyma was isolated and analyzed for percentage positive tissue area and mean intensity of positive pixels of expression for each protein. All 44 donors were available for analysis of COL1A1, COL6A2, FBLN2, LTBP4, LUM, DCN, VCAN, AND ELN, while for COL6A1, COL14A1, and FBLN5 43 donors were available. For each protein, regression coefficients (± 95% CI) were obtained following linear regression and a p value of <0.05 was considered significant. Non-transformed variables are plotted first, followed by log transformed variables. The differences in ECM and ECM-associated proteins in COPD tissue were compared to control in terms of A-C) percentage area and D-F) mean intensity of positively stained pixels. Dark blue-colored bars in light blue colored boxes highlight significant differences. ECM: extracellular matrix; SEO-COPD: Severe early-onset COPD patients; FFPE: formalin fixed paraffin embedded; COL1A1: collagen type I α chain 1; COL6A1: collagen type VI α chain 1; COL6A2: collagen type VI α chain 2; COL14A1: collagen type XIV α chain 1; FBLN2: fibulin 2; FBLN5: fibulin 5; LTBP4: latent transforming growth factor binding protein 4; LUM: lumican; DCN: decorin; VCAN: versican; ELN: elastin.

#### 3.3.2 Differences in the airway walls; Lower proportional levels of COL1A1, FBLN2, LTBP4, DCN, and VCAN in COPD and SEO-COPD

When comparing staining of ECM and ECM-associated proteins in airway walls between COPD and controls, several proteins had proportionally lower percentage area and mean intensity of expression in COPD compared to control airway walls. In the COPD group, percentage area of DCN (p=0.045) and mean intensities of COL1A1 (p=0.016) were proportionally lower compared to the control group (**Figure 3A, D**). Furthermore, subgroup analysis demonstrated proportionally lower percentage areas of COL1A1 (p=8.3x10^-5^), FBLN2 (p=0.014), and LTBP4 (p=0.016) and mean intensities of COL1A1 (p=0.013), LTBP4 (p=0.020), and VCAN (p=0.031) in the airway walls of the SEO-COPD donors compared to matched controls (**Figure 3B, E**). Similar to the parenchyma, no differences were observed when comparing moderate COPD donors to their matched controls (**Figure 3 C, F**). The percentage area and mean intensity of expression of certain ECM proteins was significantly associated with age as also demonstrated in our previous publication (12), however this did not affect the differences between COPD and control because these cohorts were age-matched (data not shown).

**Figure 3:**
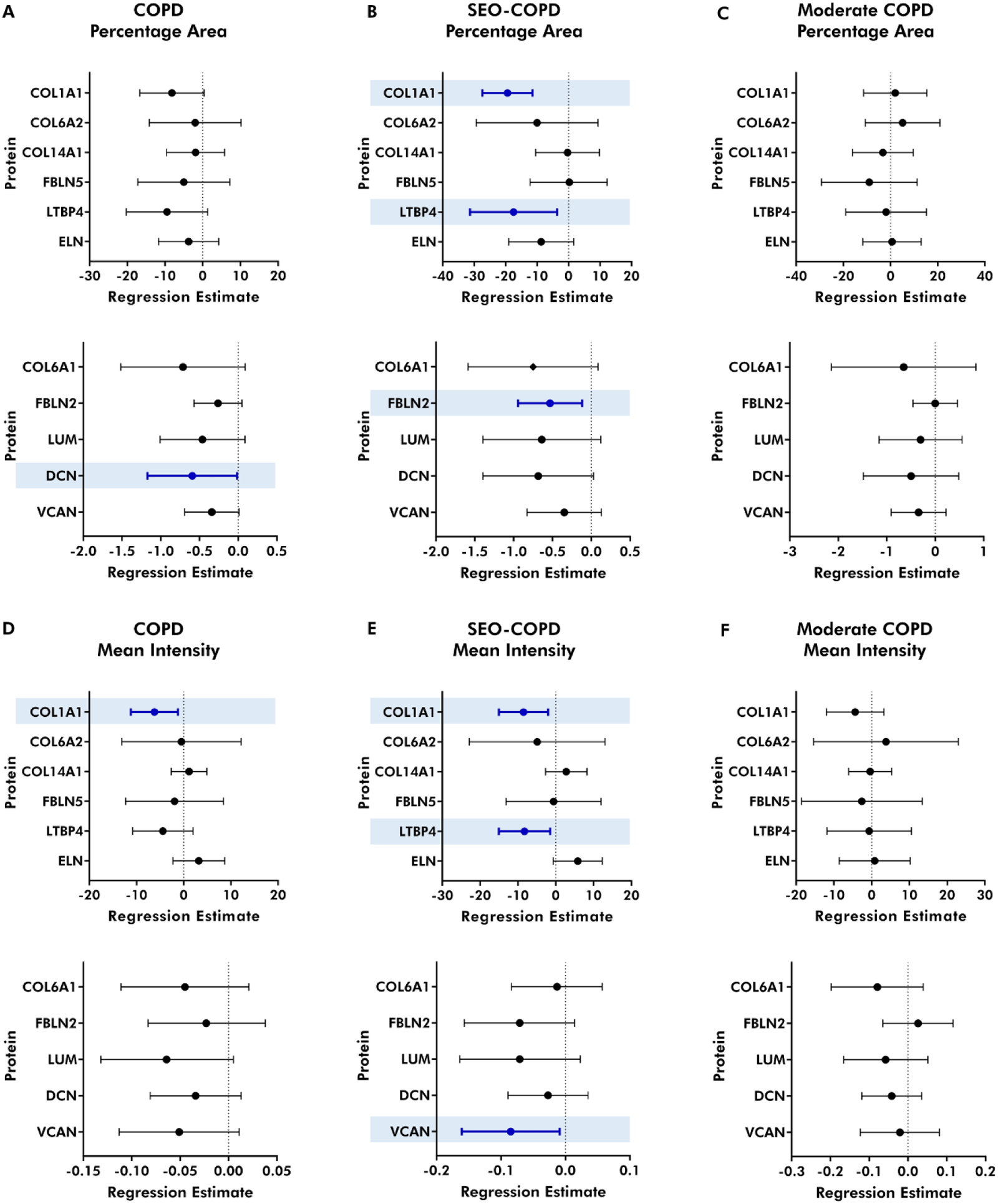
Forest plots of regression estimates for percentage area and mean intensity of ECM and ECM-associated proteins in the airway walls of COPD, SEO-COPD, and moderate COPD compared to their respective control groups. ECM proteins present in FFPE tissue sections from controls (n=18), SEO-COPD (n=12), and moderate COPD (n=14) were detected using immunohistochemistry. Airway walls were isolated and analyzed for percentage area and mean intensity of expression for each protein. The number of airway walls available for the analysis for each protein were COL1A1 (n= 155), COL6A1 (n=150), COL6A2 (n=149), COL14A1 (n=152), FBLN2 (n=158), FBLN5 (n=173), LTBP4 (n=163), LUM (n=158), DCN (n=156), VCAN (n=165), and ELN (n=158). For each protein, regression coefficients (± 95% CI) were obtained following linear regression and a p value of <0.05 was considered significant. Non-transformed variables are plotted first, followed by log transformed variables. The differences in ECM and ECM-associated proteins in COPD tissue were compared to control in terms of A-C) percentage area and D-F) mean intensity of positively stained pixels. Dark blue-colored bars in light blue colored boxes highlight significant differences. ECM: extracellular matrix; SEO-COPD: Severe early-onset COPD patients; FFPE: formalin fixed paraffin embedded; COL1A1: collagen type I α chain 1; COL6A1: collagen type VI α chain 1; COL6A2: collagen type VI α chain 2; COL14A1: collagen type XIV α chain 1; FBLN2: fibulin 2; FBLN5: fibulin 5; LTBP4: latent transforming growth factor binding protein 4; LUM: lumican; DCN: decorin; VCAN: versican; ELN: elastin.

### 3.4 Differences in the levels of ECM and ECM-associated proteins in the parenchyma and airway walls are associated with FEV1

Correlations between percentage area and mean intensity of ECM and ECM-associated proteins with lung function (FEV_1_%pred) were evaluated to investigate any relationships between proportionally lower ECM protein levels with lung function. In the parenchyma and airway walls, percentage area and mean intensity of various proteins including COL1A1, COL6A1, FBLN2, LTBP4, LUM, and VCAN were associated with FEV_1_%pred when all donors were included **(Table S1)**. On examining FEV_1_ correlations with ECM protein levels exclusively in COPD donors, percentage area and mean intensity of COL1A1 and FBLN5 in the airway walls were associated with FEV_1_ measurements **(Table S1)**. In control donors alone, percentage area and/or mean intensity of FBLN2, DCN, VCAN, and ELN were associated with FEV_1_ **(Table S1)**.

### 3.5 Identification of novel parenchymal and airway wall ECM signatures for COPD and COPD subgroups using percentage area of protein expression

Having analyzed the localization, distribution, and degree of expression of each protein individually we were also interested to explore whether there were any multi-component identifying patterns within our data set that reflected the ECM differences related to the COPD status of the patients. To explore these possible patterns, we used PCA to identify unique groupings of the proteins, that initially examined the percentage area positive staining for each protein. PCA analysis resulted in four and six components in the parenchyma and airway wall respectively, each consisting of a different set of ECM proteins, to explain at least 80% of the total variance (as seen in the scree plot) (**Figure 4A**, 4B).

**Figure 4:**
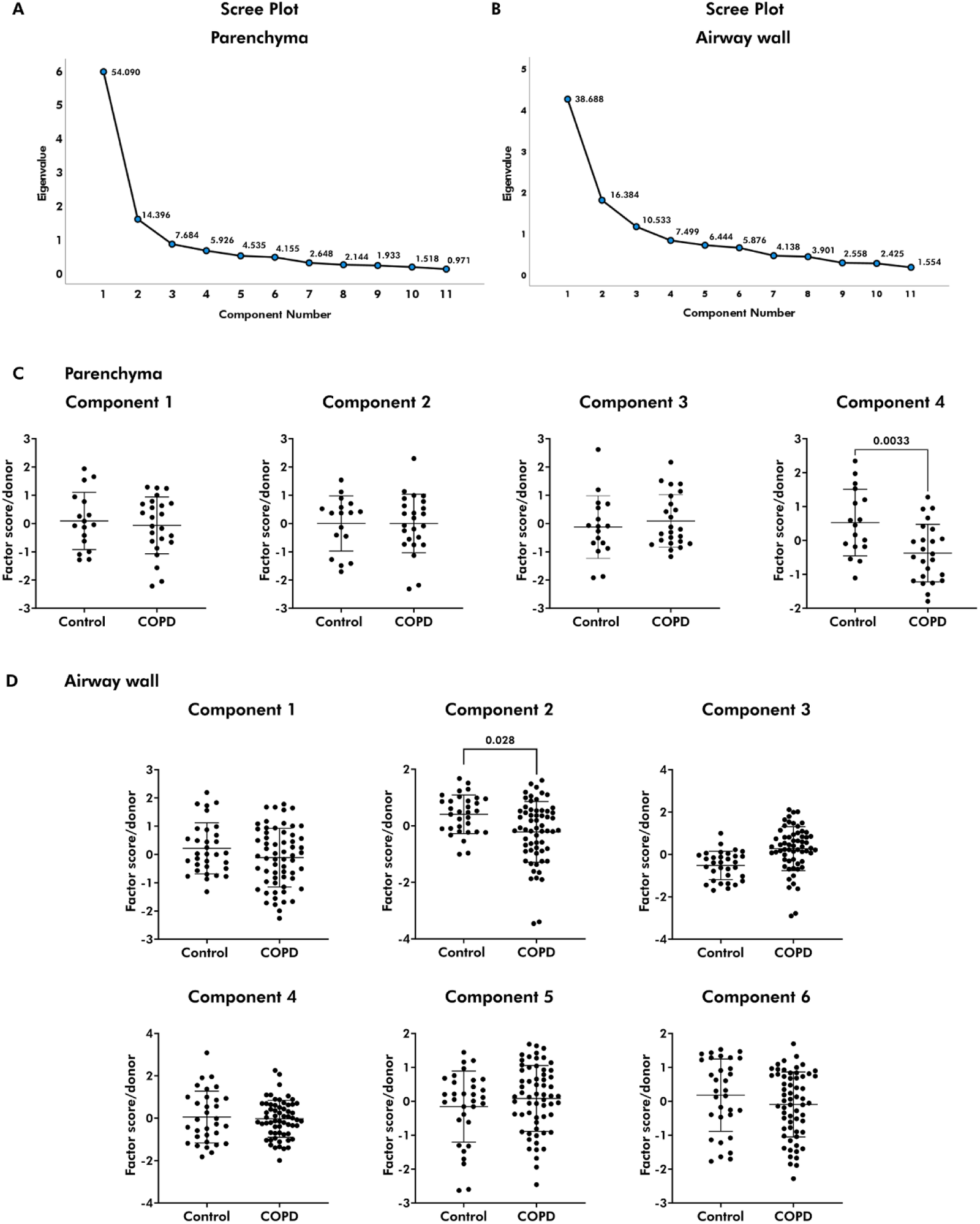
Identifying unique ECM signatures for COPD using protein expression in terms of percentage area. Principal component analysis (PCA) was performed for percentage area of proteins in parenchyma and airway walls. PCA eliminates an entire donor or airway in the absence of a measurement for even one protein out of the eleven, leaving fewer donors for parenchyma (n=41) and airway walls (n=93) in these analyzes. (A) Four and B) Six components respectively explained at least 80% of the total variance in parenchyma and airway wall as shown in the scree plot. The component scores obtained following PCA were compared using linear and linear mixed models to investigate the differences between the control and COPD groups and the mean ± SD has been plotted. C) In the parenchymal region, the patterns in proteins between COPD and control were different in component four (p=0.003). D) Differences in the patterns of proteins between control and COPD donors were noted in component two (p=0.028) in the airway walls.

When comparing the component scores for each component between COPD and control in the parenchyma, we identified a significantly lower score for the fourth component in COPD compared to controls, indicating that the combination of ECM proteins in this component reflects a parenchymal ECM signature for COPD (**Figure 4C**). Percentage area of LUM, LTBP4, DCN, COL6A1, and ELN were the highest contributors of variance in this component (**Table 3**). Subsequent subgroup analysis showed that this component was also significantly lower in SEO compared to control **(Figure S1A),** supporting our earlier observations of the differences in the parenchymal regions being mainly in the SEO-COPD patients. Thus, these 5 proteins together provide a unique ECM signature for differences in COPD and SEO-COPD parenchyma as compared to control.

In the airway walls, component scores of the COPD cohort were observed to be lower in the second component (**Figure 4D**). Percentage area of COL6A1, DCN, FBLN5, VCAN, and COL14A1 were the highest contributors to component two, thereby uniquely describing COPD airway walls (**Table 4**). Subgroup analysis indicated that the moderate group drove the differences observed in the second component **(Figure S1B)**. Additionally, SEO-COPD donors had higher scores in component three compared to controls **(Figure S1B)**. Thus COL6A1, DCN, FBLN5, VCAN and COL14A1 form a unique ECM signature for COPD airway walls, and the combined pattern of COL1A1, FBLN5, and COL14A1 describes the difference between moderate and SEO-COPD.

While these signatures are based on measurements of percentage area, mean intensities were also used to identify unique ECM signatures **(supplementary data Figure S2 and Table S2, S3)** for COPD status in parenchyma and airway walls. Mean intensities of LUM, COL6A1, LTBP4, COL6A2, and VCAN together formed an ECM signature for COPD parenchyma, while COL1A1, FBLN5, and COL6A2 described COPD airways.

**Table 3:** Rotated component matrix for principal component analysis of percentage tissue areas positive of ECM and ECM-associated proteins in the parenchyma. Four components explained at least 80% of the total variance. The loadings of each ECM and ECM-associated protein as obtained as a result of the principal component analysis are tabulated below. The loadings represent the correlations between the proteins and the component.

| Rotated Component Matrix - Parenchyma |  |  |  |  |
| --- | --- | --- | --- | --- |
|  | 1 | 2 | 3 | 4% |
| VCAN | 0.814 | 0.308 |  |  |
| FBLN2 | 0.802 |  | 0.376 |  |
| ELN | 0.751 |  | 0.399 | 0.308 |
| COL14A1 |  | 0.918 |  |  |
| FBLN5 | 0.439 | 0.791 |  |  |
| COL6A1 | 0.373 | 0.662 | 0.311 | 0.333 |
| LTBP4 | 0.532 | 0.533 |  | 0.468 |
| DCN | 0.345 | 0.489 | 0.483 | 0.387 |
| COL1A1 | 0.398 |  | 0.839 |  |
| COL6A2 |  | 0.472 | 0.637 |  |
| LUM |  |  |  | 0.917 |

%p <0.005 and indicates the comparison between COPD donors and controls. Only loadings > |0.3| are shown.

**Table 4:** Rotated component matrix for principal component analysis of percentage tissue areas positive of ECM and ECM-associated proteins in the airway walls. Six components explained at least 80% of the total variance. The loadings of each ECM and ECM-associated protein as obtained as a result of the principal component analysis are tabulated below. The loadings represent the correlations between the proteins and the component.

| Rotated Component Matrix- Airway walls |  |  |  |  |  |  |
| --- | --- | --- | --- | --- | --- | --- |
|  | 1 | 2% | 3%% | 4 | 5 | 6 |
| ELN | 0.918 |  |  |  |  |  |
| FBLN2 | 0.762 |  |  |  | 0.400 |  |
| VCAN | 0.633 | 0.482 |  | 0.300 |  |  |
| COL6A1 |  | 0.873 |  |  |  |  |
| DCN | 0.496 | 0.663 |  |  |  |  |
| COL1A1 |  |  | -0.823 |  |  |  |
| FBLN5 |  | 0.557 | 0.710 |  |  |  |
| LTBP4 | 0.471 |  |  | 0.788 |  |  |
| COL14A1 |  | 0.336 | 0.367 | 0.698 |  |  |
| COL6A2 |  |  |  |  | 0.893 |  |
| LUM |  |  |  |  |  | 0.977 |

%p <0.05 and indicates the comparison between COPD donors and controls. %%p<0.05 and indicates the comparison between SEO-COPD, moderate, and control donors corrected for age. Only loadings > |0.3| are shown.

### 3.6 COPD-associated differences in the lung ECM

We have summarized the differences seen in the landscape of ECM in the lung parenchyma and airway walls in COPD and the subgroups (SEO and moderate) in **Figure 5**. Only those proteins that were observed to have significant associations with COPD or its subgroups have been indicated.

**Figure 5:**
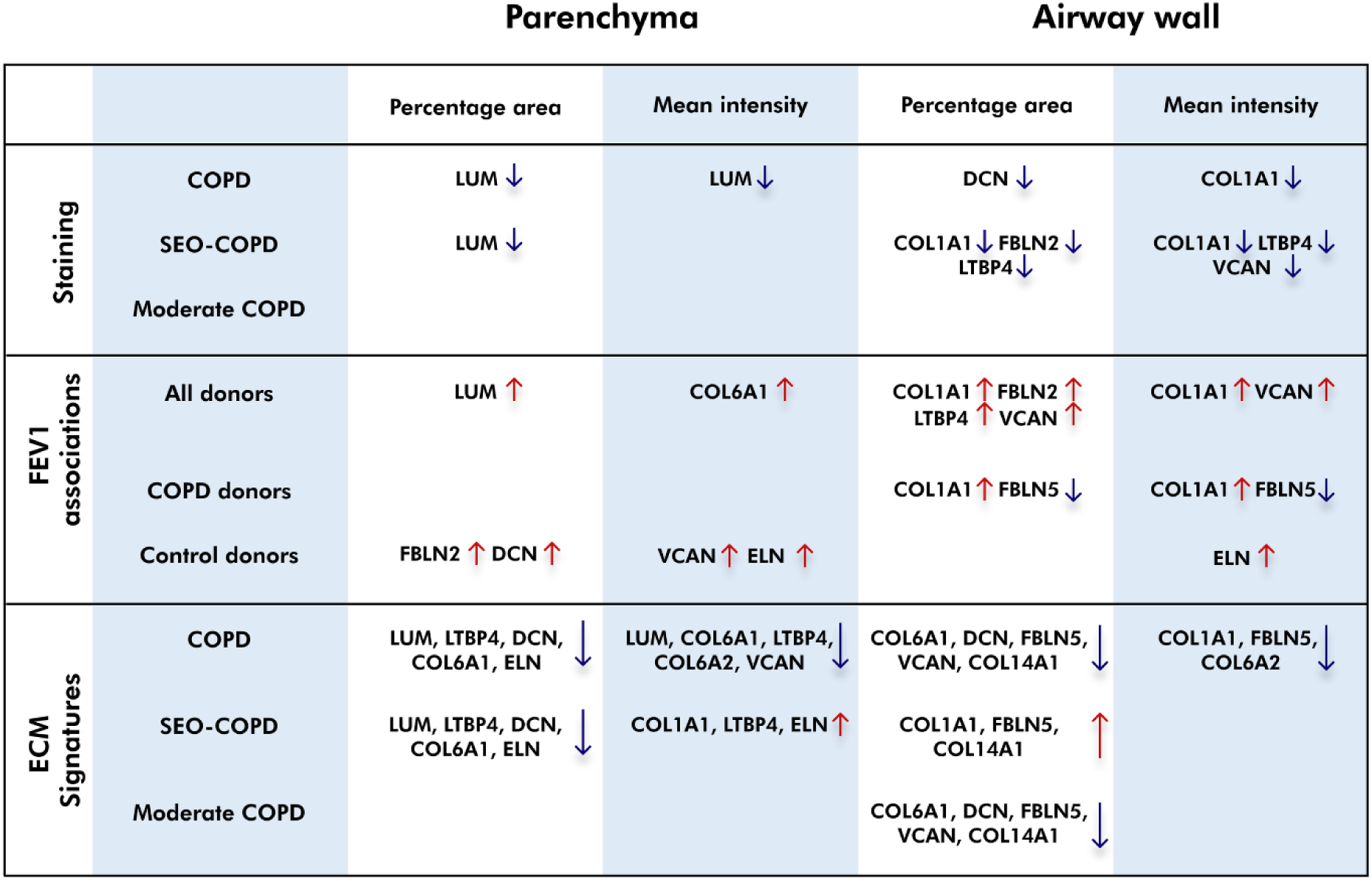
COPD-associated differences in lung ECM. ECM differences noted in the different analyzes throughout this study have been summarized here. In the staining and ECM signatures, red or blue arrows indicate higher or lower proportional levels or component scores in COPD respectively, while they denote positive or negative associations with FEV1 respectively. ECM: extracellular matrix; SEO-COPD: Severe early-onset COPD patients; COL1A1: type I collagen α chain 1; COL6A1: type VI collagen α chain 1; COL6A2: type VI collagen α chain 2; COL14A1: type XIV collagen α chain 1; FBLN2: fibulin 2; FBLN5: fibulin 5; LTBP4: latent transforming growth factor binding protein 4; LUM: lumican; DCN: decorin; VCAN: versican; ELN: elastin; FEV1: forced expiratory volume in 1 second.

## 4. DISCUSSION

Our study provides an elaborate investigation of eleven ECM and ECM-associated proteins in COPD lung tissue. Important novelties of this study are the combination of the comparison between airway walls and lung parenchyma, the inclusion of different COPD subgroups, characterization of proteins that have not yet been studied in COPD lung tissue, and the investigation of mean intensity, in addition to percentage tissue area of expression of each protein. Overall, the SEO-COPD group showed the most ECM differences in composition compared to controls. Notably, we generated unique ECM signatures that described moderate and SEO-COPD independently, potentially reflecting differing tissue remodeling processes that are part of the disease pathogenesis in these COPD subtypes.

ECM dysregulation has been postulated as a common hallmark of aging and COPD (8). Previously our group identified specific profiles of age-associated genes and a negative interaction between age and presence of COPD for several ECM-related genes, indicating a different association of age with COPD and control (20). Our follow-up study focused on ECM differences with normal aging and revealed higher expression of COL1A1, COL6A1, COL6A2, FBLN2, LTBP4, and LUM at both gene and protein levels in aging lungs of control patients with normal lung function and no history of COPD, pulmonary fibrosis or asthma (12). Additionally, immunohistochemical analysis showed higher levels of COL6A2 in the airway walls and COL6A2 and COL1A1 in the parenchyma with aging lungs in control subjects. This age-related ECM profile in control patients did not overlap with the ECM signature reported for COPD in the present study, where we observed proportionally lower ECM and ECM-associated protein percentage area and/or mean intensity in COPD. This suggests that the observed ECM profile in COPD is different from age-related ECM differences in non-COPD controls, and can be either driven by the pathobiology of COPD or represent a form of abnormal aging in COPD. Notably, parenchyma in both aging and COPD is affected by emphysema, however, changes in the airway walls structures are less comparable between the two as airway thickening is not as apparent in aging lungs.

ECM differences in COPD have long been an area of interest (21). However, results from different studies are often conflicting. In the current study, we identified more COPD-associated differences in the airway walls compared to the parenchyma. It is important to note that, as a result of emphysematous loss of the lung parenchyma, tissue obtained from COPD patients in the more severe stages only allows us to examine the parenchyma and airways that are still remaining in the lung. It is quite likely that the ECM composition of the lost tissue was also aberrant, however, this cannot be characterized *ex vivo*. Moreover, we report proportional differences in the ECM content in the airway walls in COPD compared to control tissue. Proportional differences do not necessarily indicate that there is less total ECM content in the airway wall, but rather that the proportion of the different ECM components is changing. Thus, there may be an absolute increase in other ECM proteins in the airways, due to the fibrotic nature of the changes observed in COPD, that have not captured in this study. It is thus important to assess more ECM proteins, such as collagen III as a relative increase in COL3 over COL1 has been previously demonstrated in COPD tissue (24). In parenchyma and airway walls alike, protein levels did not differ between the moderate COPD group and their matched controls. The ECM profiles in the moderate COPD patients, who were older compared to SEO-COPD patients, also did not resemble the differences in ECM as noted previously in aging lungs. It is clear from the present study that the pathogenesis of moderate and SEO-COPD in terms of ECM landscape is different and capturing data in SEO-COPD patients earlier would help understand the similarities in disease mechanisms between the two subgroups. Unfortunately, SEO-COPD patients are often diagnosed at the stage that their symptoms are quite severe and the tissue collected is collected at the time of lung transplantation, thus early stage data is scarce.

Collagens are the most abundant proteins in the lung ECM. Fibrillar collagens (type I and III) are crucial in maintaining the structural integrity and organization of the lung tissue by forming 3D networks and providing mechanical strength (22). Less fractional area and volume fraction of collagen type I in COPD lungs has previously been reported in small airways by Annoni *et al*. (23) and bronchiolar tissue by Hogg *et al*. (24) respectively. These findings align with the lower proportion of COL1A1 in airway walls of COPD patients observed in our study. However, neither of these previous studies compared the differences between COPD severities, while we observed that SEO-COPD donors dominantly contribute to the lower proportion of COL1A1 in COPD within the airway wall. In parenchyma, Eurlings *et al*. (25) demonstrated higher percentage area of collagen (type I/II/III) in the remaining alveolar walls of COPD subjects which increased with disease severity, whereas we did not observe differences in the proportion of COL1A1 in the parenchyma. Eurlings *et al.* examined older COPD stage IV patients, compared to the relatively younger SEO-COPD donors in our study, providing a possible explanation for the inconsistent findings between both studies.

Another category of ECM proteins are proteoglycans and we have investigated LUM, DCN, and VCAN. LUM, DCN, and VCAN play a role in various cellular functions, bind to growth factors and chemokines, and regulate fibrillogenesis (26–30). The fractional area of LUM in COPD lung tissue compared to controls has been previously reported by Annoni *et al*. (23) who did not observe any differences between COPD donors and controls. In contrast to Annoni’s study, we showed lower percentage area of LUM in the parenchymal region of COPD patients, in particular in SEO-COPD. Annoni *et al.* also reported lower VCAN fractional area in parenchyma in COPD and Merrilees *et al*. (31) observed that the alveolar walls of COPD patients showed stronger VCAN staining than control donors. In our study lower mean intensity of VCAN was observed in the airway walls of SEO-COPD patients. Consistent with our finding of lower proportional percentage area of DCN in COPD airway walls, a previous study reported lower DCN staining in the airway adventitia of severe emphysematous tissue, while only a few donors with mild emphysema displayed lower staining (15). However, in our current study we did not observe an association between DCN staining intensity and disease severity. The loss of proteoglycans can, not only directly alter cellular responses (32), but also affect the structural integrity of lung tissue due to the resulting disorganized ECM components and modified mechanical properties such as lung elasticity and alveolar stability (33).

FBLNs are calcium-binding glycoproteins that bind to other ECM aggregates and stabilize them. They bind to the basement membrane and also elastic fibers (34). In our previous study we observed a positive correlation between percentage area of FBLN2 and the mean intensity of COL1A1 in the parenchyma with age in control never-smokers (12). Interestingly, in COPD lungs, a lower percentage area of FBLN2 was accompanied by a lower percentage area and mean intensity of COL1A1 in SEO-COPD donors in airway walls. Despite both being observational studies, these results incite further investigation into the FBLN2/COL1A1 axis as a possible regulatory mechanism for the stabilization of collagen fibers around the airways. (23, 25, 35). An earlier study from our group reported higher total levels of FBLN5 in COPD tissue homogenates compared to controls using western blot, along with colocalization of FBLN5 with ELN fibers using immunohistochemical staining in lung tissue (36).

Next to assessing all ECM differences separately, it is also relevant to study the patterns in ECM differences. Together these differences form the ECM landscape in COPD and may contribute to the change in mechanical properties of the lung such as loss of elastic recoil and airflow obstruction of COPD airways. This is supported by the correlation of the expression of several of these proteins with FEV_1_ measurements. Moreover, a recent study reported softer emphysematous precision cut lung slices with decreased stiffness compared to healthy controls (37). With the help of computation network modeling, they attributed this reduced stiffness to ECM remodeling of the septal wall along with structural deterioration of the lung. In the current study, we identified ECM signatures that represent groups of proteins the characteristics of which describe COPD and the different severities of COPD. It is clear from the various results obtained in our study that LUM, COL6A1, and LTBP4 are important in describing COPD parenchyma ECM characteristics. Similarly, COL1A1, COL14A1, FBLN5, DCN, and VCAN appear essential in describing characteristic factors in ECM remodeling in the airway walls in COPD (**Figure 5**).

Collagen type I and III are major components of the lung ECM. However, several supporting ECM and ECM-associated proteins are required to maintain the structural integrity and function of collagens. Per our knowledge, expression of LTBPs in lung tissue has not been studied in the context of COPD. LTBPs not only regulate the signaling of transforming growth factor β (TGF-β), but also mediate elastogenesis (38, 39). LTBP4 binds to TGF-β1and guides its deposition into the ECM after cellular secretion. Thus, less proportional presence of LTBP4 may result in lower TGF-β1 activation and diminished elastogenesis. Another consequence of decreased TGF-β-Smad pathway activation is the attenuation of DCN in COPD lungs and hence disruption of collagen assembly (40). Moreover, abnormal loosening of collagen fibers might make ELN susceptible to destruction. A study in adult rats reported that the turnover-rate of collagens is 10-15% per day (41), however, formation of ELN fibers declines with age and is almost absent in adulthood (42). Additionally, relatively lower protein levels of factors regulating elastogenesis, such as LTBP4 and FBLN2 in the current study, potentially hamper ELN repair. In our study, however, we did not observe a difference in the percentage area or mean intensity of ELN. Several studies have shown a reduction of ELN fibers in airways and parenchyma; however, no differences were noted between different stages of COPD (23, 35). Notably, the unique signatures identified for airway walls and parenchyma suggest compartmental differences during reparative processes. Among proteoglycans, LUM was proportionally lower in the parenchyma while DCN and VCAN were lower in the airway walls. This could indicate that LUM is more closely associated with collagen assembly in the parenchyma, while DCN and VCAN are more critical in the airway walls. While most proteins appear in either the parenchymal or airway wall signature alone, COL6 appears in both. It is localized near the basement membrane region forming a meshwork between the basement membrane and the interstitial matrix. The role of COL6 in mechanical regulation of the lung has previously been postulated (43). Its presence in both signatures suggests a key role for COL6 in the overall COPD pathology. Collagen type XIV is found in regions of high mechanical strength. It is well-known that the mechanical properties of airway walls in COPD, including fibrosis and loss of elastic recoil, are altered (44). Thus, it is not surprising that we observed COL6A1, COL6A2 and COL14A1 localized in the airway smooth muscle region and these proteins appear important in our airway wall signatures. Our results suggest that COL6 and COL14A1 likely are important contributors to airway wall pathology that leads to the obstruction and ultimately the collapse of airways in COPD.

A limitation of this study is the small number of patients included in the sub-group analysis. The heterogeneity of COPD, patient inclusion criteria, and analysis methods may account for the variability in the data seen among these studies. Despite these limitations we reported robust findings. Our results not only suggest novel target pathways for developing therapies for alleviating COPD symptoms but also are a reminder that disease pathogenesis is not driven and characterized by a single protein but rather by a complex interactions between groups of proteins.

In conclusion, we identified COPD-associated differences in the lung ECM composition. The differences in the ECM landscape may affect the integrity of the structure and function of the lung tissue compartments. Our study has identified proteins, including LUM in the parenchyma and COL1A1, LTBP4, FBLN2, and VCAN in the airway walls that have not previously been associated with the SEO-COPD subtypes. Moreover, we report unique ECM signatures for parenchyma and airway walls associated with disease and COPD severity. The identified proteins may contribute to the establishment and maintenance of disease pathology in SEO-COPD patients and thus provides leads for future studies.

## 5. GRANTS

MMJ acknowledges support from the Graduate School of Medical Sciences, University of Groningen. NB is funded by the PPP-allowance by Health Holland (LSH) and the UMCG. This collaboration project is co-financed by the Ministry of Economic Affairs and Climate Policy by means of the PPP-allowance made available by the Top Sector Life Sciences & Health to stimulate public-private partnerships. MLKN receives support from Abel Tasman Talent Program Fellowship in association with the Healthy Aging Alliance from the UMCG. RRW acknowledges support from Lung Foundation Netherlands (Longfonds 5.2.21.006). CAB receives from Nederlandse Organisatie voor Wetenschappelijk Onderzoek (NWO) (Aspasia 015.015.044). JKB acknowledges support from the NWO (Aspasia 015.013.010).

