## Supplemental material for "The lung extracellular matrix protein landscape in severe early-onset and moderate chronic obstructive pulmonary disease"

***Identification of novel parenchymal and airway wall ECM signatures for COPD and COPD subgroups using mean intensity of protein expression***

In addition to identifying ECM signatures based on percentage area of protein expression, we were interested in identifying patterns within our data using mean intensity of protein expression as well. The data were described by five and six components in the parenchyma and airway walls respectively, to explain at least 80% of the total variance as seen in the scree plot **(Figure S2A, S2B).**

In the parenchyma, we identified significantly lower scores of the COPD cohort in the second component in comparison to the controls while comparing the factor scores for each principal component. **(Figure S2C)**. Mean intensity of LUM, COL6A1, LTBP4, COL6A2, and VCAN were the highest contributors of variance in the second component **(Table S2)**. Subsequently on analyzing the subgroups, that is on segregation of the SEO and moderate COPD donors **(Figure S2E)**, an ECM signature including COL1A1, ELN, and LTBP4 **(Table S2)** could describe differences between control and SEO-COPD donors and SEO-COPD and moderate COPD donors.

In the airway walls, component three and component six with lower factor scores were observed to have groups of proteins that uniquely described COPD **(Figure S2D).** In the airway walls, component three was composed of COL1A1, COL6A2, and FBLN5 that could describe the difference between COPD and control airway walls **(Table S3).** Subgroup analysis yielded no further significant differences between the different groups. Therefore, for the airway walls the mean intensities of COL1A1, COL6A2, and FBLN5 was considered as the unique ECM signature for COPD airway walls.

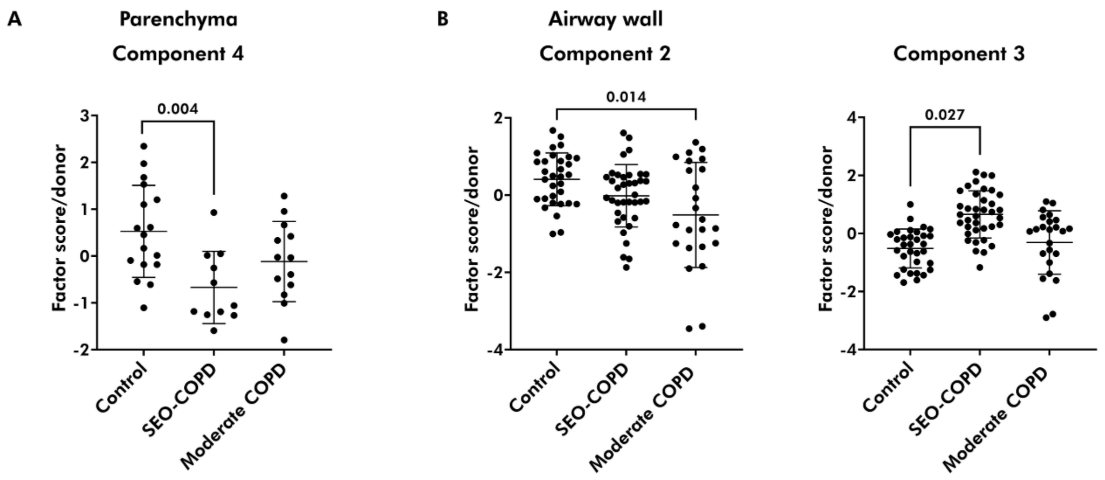


**Figure S1: Subgroup analysis using factor scores of SEO and moderate COPD donors compared to controls.** For subgroup analysis, factor scores obtained following principal component analysis (PCA) were compared using linear and linear mixed regression and the mean ± SD has been plotted. PCA eliminates an entire donor or airway in the absence of a measurement for even one protein out of the eleven, leaving fewer donors for parenchyma (n=41) and airway walls (n=93) in these analyses. These comparisons revealed that A) The SEO-COPD donors drove the overall difference in component four in parenchyma (p= 0.004). B) In the airway walls, moderate COPD donors majorly contributed to the difference between controls and COPD in component two (p= 0.014), while SEO-COPD donors did so in component three (p= 0.027). SEO-COPD: Severe early-onset COPD patients.

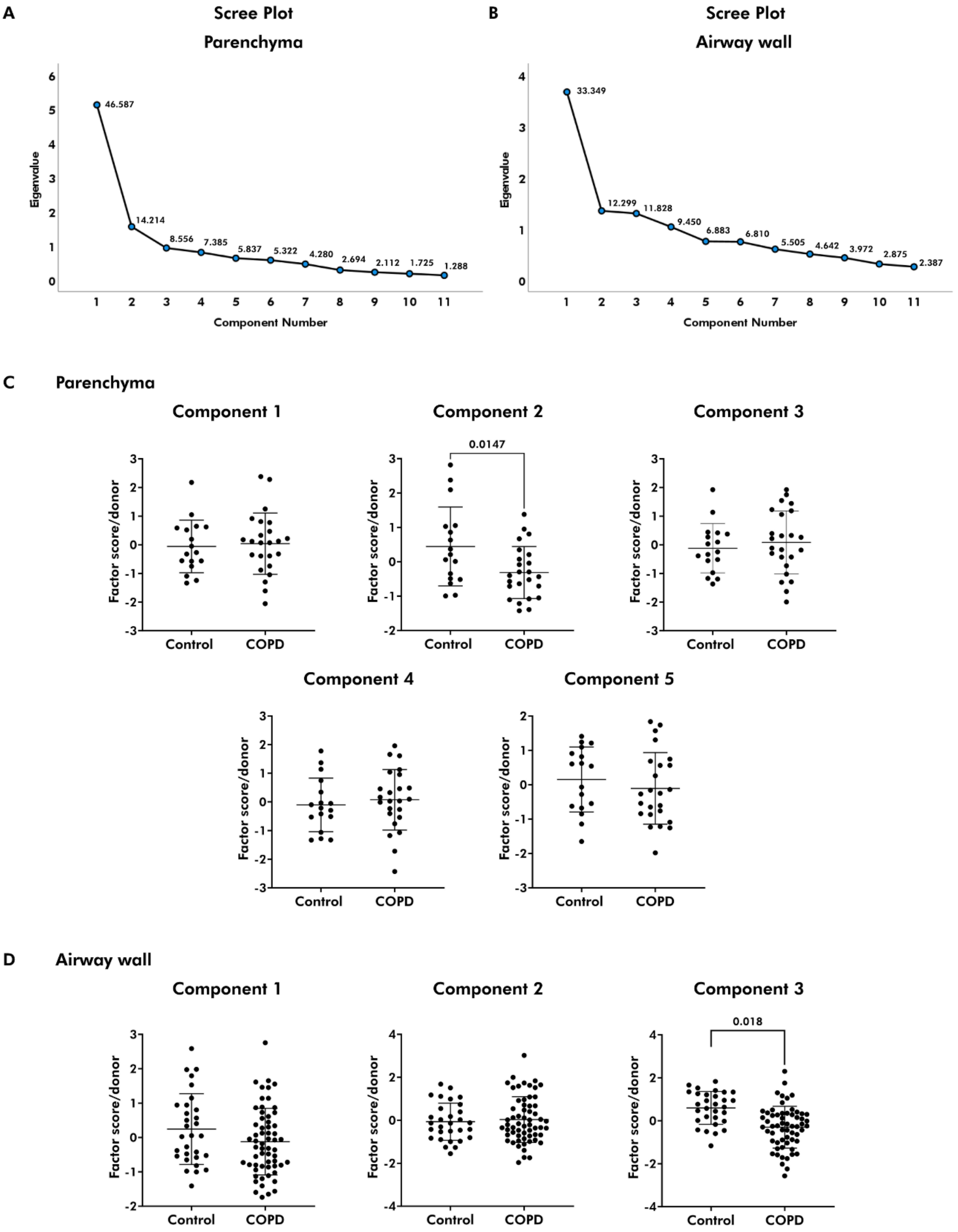
A
Scree Plot
Parenchyma
Eigenvalue
Component Number
46.587
14.214
8.556
7.385
5.837
5.322
4.280
2.694
2.112
1.725
1.288
B
Scree Plot
Airway wall
Eigenvalue
Component Number
33.349
12.299
11.828
9.450
6.883
6.810
5.505
4.642
3.972
2.875
2.387
C
Parenchyma
Component 1
Component 2
0.0147
Component 3
Component 4
Component 5
Factor score/donor
Control
COPD
D
Airway wall
Component 1
Component 2
Component 3
0.018
Factor score/donor
Control
COPD

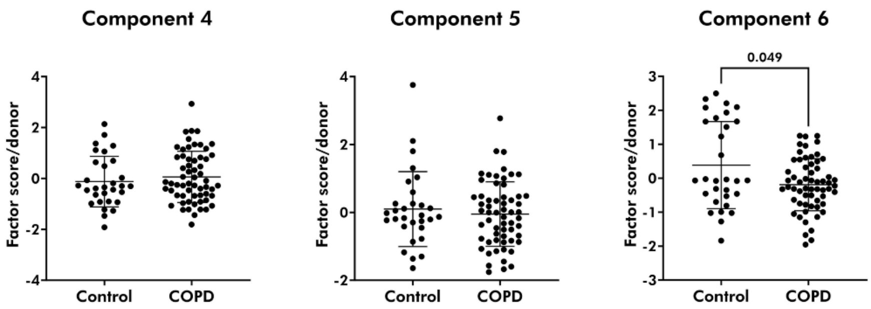


E

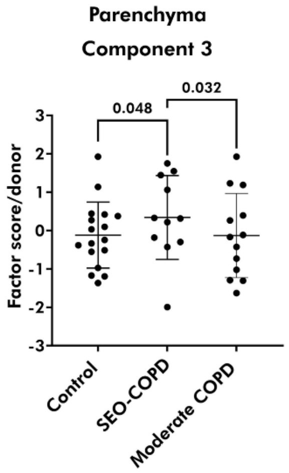


***Figure S2: Identifying unique ECM signatures for COPD using protein expression in terms of mean intensity.*** Principal component analysis (PCA) was performed for mean intensities of proteins in parenchyma and airway walls. PCA eliminates an entire donor or airway in the absence of a measurement for even one protein out of the eleven, leaving fewer donors for parenchyma (n=41) and airway walls (n=93) in these analyses. A) Five and B) Six components explained at least 80% of the total variance in parenchyma and airway walls as shown in the scree plot. The component scores obtained following PCA were compared linear regression to investigate the contribution of each donor in the control and COPD groups and the mean ± SD has been plotted. C) In the parenchymal region, the patterns in proteins between COPD and control were different in component two (p=0.0147). D) Differences in the patterns of proteins between control and COPD donors were noted in component three (p= 0.018) and six (p=0.0496) in the airway walls. E) Subgroup analysis revealed that SEO-COPD donors had higher contribution compared to both moderate COPD (0.032) and control (p=0.048) donors in component three. SEO-COPD: Severe early-onset COPD patients.

***Table S1:* Associations of ECM and ECM-associated proteins with lung function measurements.** Correlations between percentage area and mean intensity of each protein in the parenchyma and airway walls with $FEV_1$%pred were calculated for all donors and for COPD donors exclusively. Significant associations ($p<0.05$) have been depicted on a scale from white to red for positive and white to blue for negative associations.

| Protein | Parenchyma | | Airway wall | |
|---|---|---|---|---|
| | Percentage Area | Mean intensity | Percentage Area | Mean intensity |
| **All donors** | | | | |
| COL1A1 | | | | |
| COL6A1 | | | | |
| COL6A2 | | | | |
| COL14A1 | | | | |
| FBLN2 | | | | |
| FBLN5 | | | | |
| LTBP4 | | | | |
| LUM | | | | |
| DCN | | | | |
| VCAN | | | | |
| ELN | | | | |
| **COPD donors** | | | | |
| COL1A1 | | | | |
| COL6A1 | | | | |
| COL6A2 | | | | |
| COL14A1 | | | | |
| FBLN2 | | | | |
| FBLN5 | | | | |
| LTBP4 | | | | |
| LUM | | | | |
| DCN | | | | |
| VCAN | | | | |
| ELN | | | | |
| **Control donors** | | | | |
| COL1A1 | | | | |
| COL6A1 | | | | |
| COL6A2 | | | | |
| COL14A1 | | | | |
| FBLN2 | | | | |
| FBLN5 | | | | |
| LTBP4 | | | | |
| LUM | | | | |

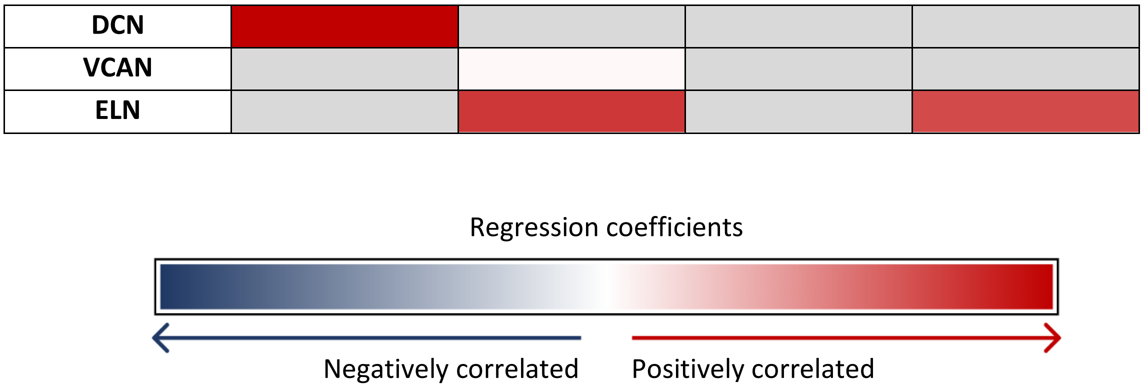


FEV$_1$%pred measurements were missing for two control (moderate COPD matched) donors. COL1A1: type I collagen α chain 1; COL6A1: type VI collagen α chain 1; COL6A2: type VI collagen α chain 2; COL14A1: type XIV collagen α chain 1; FBLN2: fibulin 2; FBLN5: fibulin 5; LTBP4: latent transforming growth factor binding protein 4 ; LUM: lumican; DCN: decorin; VCAN: versican; ELN: elastin.

**Table S2: Rotated component matrix for principal component analysis of mean intensities of ECM and ECM-associated proteins in the parenchyma.** Five components explained at least 80% of the total variance. The loadings of each ECM and ECM-associated protein as obtained as a result of the factor analysis. The loadings represent the correlations between the proteins and the component.

| Rotated Component Matrix - Parenchyma | | | | | |
|---|---|---|---|---|---|
| | **1** | **2[%]** | **3[%%]** | **4** | **5** |
| **VCAN** | 0.845 | 0.348 | | | |
| **FBLN5** | 0.806 | | | 0.362 | |
| **FBLN2** | 0.706 | | | | 0.328 |
| **ELN** | 0.657 | | 0.588 | | |
| **COL6A1** | 0.551 | 0.540 | | 0.306 | 0.394 |
| **COL6A2** | 0.457 | 0.453 | | | 0.331 |
| **LUM** | | 0.932 | | | |
| **COL1A1** | | | 0.879 | | |
| **COL14A1** | | | | 0.941 | |
| **LTBP4** | | 0.454 | 0.452 | 0.536 | |
| **DCN** | | | | | 0.891 |

[%]p <0.05 and indicates the comparison between COPD donors and controls. [%%]p<0.05 and indicates the comparison between SEO-COPD, moderate, and control donors corrected for age. Only loadings > |0.3| are shown.

**Table S3: Rotated component matrix for principal component analysis of mean intensities of ECM and ECM-associated proteins in the airway walls.** Six components explained at least 80% of the total variance. The loadings of each ECM and ECM-associated protein as obtained as a result of the factor analysis. The loadings represent the correlations between the proteins and the component.

| Rotated Component Matrix – Airway walls | | | | | | |
|---|---|---|---|---|---|---|
| | **1** | **2** | **3[%]** | **4** | **5** | **6[%]** |
| **VCAN** | 0.829 | | | | | |
| **FBLN2** | 0.803 | | | | | |
| **ELN** | 0.773 | | | | | |
| **COL14A1** | | 0.908 | | | | |
| **DCN** | 0.504 | 0.540 | | | | |
| **COL1A1** | | | 0.857 | | | |
| **FBLN5** | 0.502 | 0.324 | -0.595 | | | |
| **COL6A2** | | | -0.304 | 0.803 | | |
| **LTBP4** | 0.380 | | | 0.618 | | |
| **COL6A1** | | | | | 0.961 | |
| **LUM** | | | | | | 0.970 |

[%]p <0.05 and indicates the comparison between COPD donors and controls. Only loadings > |0.3| are shown.
